# Depolymerases unmask a conserved receptor to enable phage control of capsulated *Acinetobacter baumannii*

**DOI:** 10.64898/2026.11.16.766953

**Authors:** Leo Kan, Laura Avellaneda-Franco, Dinesh Subedi, Denis Korneev, Ella Gillis, Isabelle Magnin-Bougma, Mark B. Carascal, Shankar Raj Devkota, Surendra Kunwar, Ram Prasad Bhusal, Ruth M. Hall, Johanna J. Kenyon, Fernando Gordillo Altamirano, Anton Y. Peleg, Jeremy J. Barr

**Affiliations:** School of Biological Sciences, Monash University, Clayton, Victoria, Australia; Centre to Impact AMR, Monash University, Clayton, Victoria, Australia; Ramaciotti Centre for Cryo-Electron Microscopy, Monash University, Clayton, Victoria, Australia; Clinical and Translational Research Institute, The Medical City, Pasig, Philippines; Department of Biochemistry & Molecular Biology, Biomedicine Discovery Institute, Monash University, Clayton, Victoria, Australia; School of Chemical Engineering, Adelaide University, Adelaide, South Australia, Australia; School of Life & Environmental Sciences, The University of Sydney, New South Wales, Australia; School of Pharmacy & Medical Sciences, Griffith University, Gold Coast, Queensland, Australia; Department of Infectious Diseases, The Alfred Hospital and School of Translational Medicine, Monash University, Melbourne, Victoria, Australia; Infection Program, Monash Biomedicine Discovery Institute, Department of Microbiology, Monash University, Clayton, Victoria, Australia

## Abstract

We characterized bacteriophage øAbs1 that infects clinical *Acinetobacter baumannii* isolates via the highly prevalent and conserved outer-membrane protein receptor, OprC. However, øAbs1 is unable to infect capsulated *A. baumannii* strains as the capsule prevents phage access to its underlying receptor. To circumvent this, we combined øAbs1 with recombinant depolymerases targeting the major *A. baumannii* capsule types. In a murine bacteremia model, this combination significantly reduced bacterial burden caused by diverse, capsulated *A. baumannii* strains.

---

*Acinetobacter baumannii* is a critical-priority pathogen whose extensive drug resistance and virulence underscore the urgent need for new infection control and treatment strategies^1,2^. A defining characteristic of *A. baumannii* is its capsule^3^, composed of structurally diverse capsular polysaccharides (CPS)^4^ that form a protective barrier of variable composition and thickness. The genes responsible for CPS biosynthesis and transport are encoded within a defined K locus (KL) in the *A. baumannii* genome^5^, providing a genetic basis for characterizing the extensive diversity of CPS types. Furthermore, the capsule plays pivotal roles in bacterial virulence^6^ and immune evasion^7^. Recently, phage therapy has emerged as a promising alternative treatment for multidrug-resistant *A. baumannii,* but its clinical potential is constrained by the narrow host range of most phages, which typically recognize and infect only a specific KL type^8,9^. With more than 400 *A. baumannii* KL types predicted^10^, developing broad-host range phage products that overcome capsule-mediated barriers against this pathogen is an urgent priority.

Here, we present an investigation of the bacteriophage øAbs1, a virulent Myovirus-like phage (**Fig. 1a**) that was originally isolated using the clinical *A. baumannii* strain LUH5538^11^ (KL83). Whole genome sequencing of øAbs1 (102,398 bp) found no integration or virulence associated genes and classified the phage within the genus *Saclayvirus* (family *Luriaviridae*, class *Caudoviricetes*) (Extended Data Fig. 1a). Growth curves of øAbs1 with its host strain revealed strong suppression of bacterial growth for up to 8h, followed by highly variable bacterial suppression, with some replicates exhibiting bacterial clearance whereas others showed bacterial regrowth, consistent with the emergence of phage-resistant variants (**Fig. 1b**). One-step growth curve showed øAbs1 had a burst size of 3.5 virions per cell and a latent period of 25 min (**Fig. 1c**). Next, to identify the receptor of øAbs1, we attempted to generate øAbs1-resistant mutants using its host of isolation (LUH5538). However, resistance was initially unstable, as phage-resistant mutants spontaneously reverted to their wild-type, phage-sensitive variants upon subculture. Eventually, stable phage-resistant mutants were isolated and sequenced. However, short-read sequencing at high depth detected no mutations relative to the parental host genome despite the phage-resistant phenotype. Given the capsule’s central role in phage infectivity in this pathogen^9^, we quantified the production of CPS between the LUH5538 wild-type and resistant strain, revealing a two-fold increase in CPS associated with øAbs1 phage-resistance (**Fig. 1d**). Based on this, we hypothesized that øAbs1 may infect *A. baumannii* via a non-capsule receptor, with the overproduction of CPS providing a transient resistance mechanism that would physically impede phage adsorption to an underlying surface receptor.

**Figure 1.**
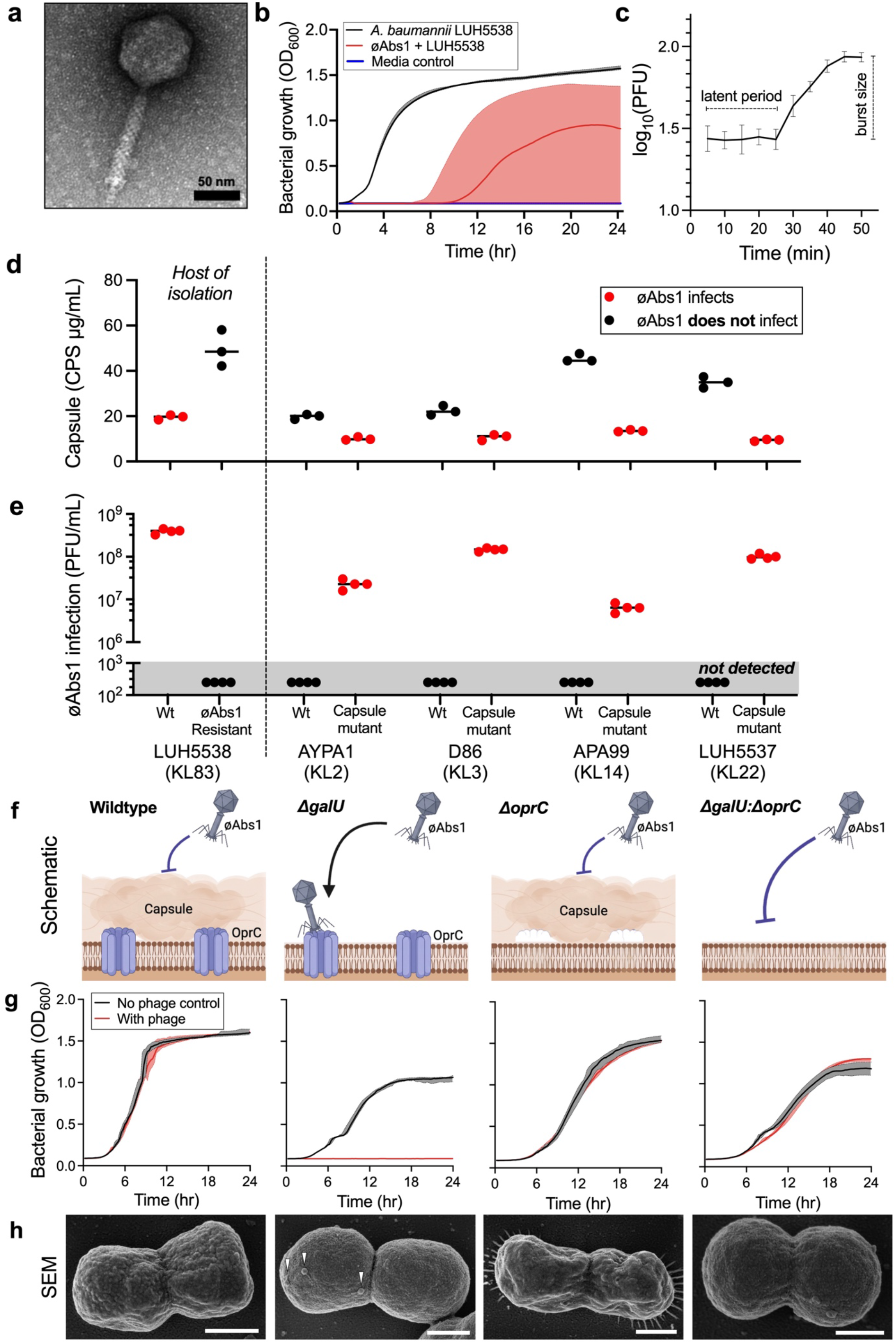
**a.** Transmission electron microscopy (TEM) image of øAbs1 (scale bar = 50 nm). **b.** Growth curves of *Acinetobacter baumannii* strain LUH5538 (black), in the presence of øAbs1 (red) at a multiplicity of infection (MOI) of 1, and media control (gray). Lines represent the median, and shaded regions indicate the 95% confidence intervals (n = 9), with the wider interval in the øAbs1-treated group reflecting greater variability among replicates. OD_600_, optical density at 600 nm. **c.** One-step growth curve of øAbs1 with host of isolation *A. baumannii* LUH5538. Data are mean ± s.d. (n = 9). **d.** Capsule quantification of wild-type and mutant *A. baumannii* strains colored by øAbs1 infectivity (red) and no infectivity (black). Individual data points are shown, line is median (n = 3). **e.** Matched øAbs1 infectivity of strains from panel d, reported in Plaque Forming Units (PFU) per mL. Individual data points are shown, line is median (n = 3), limit of detection = 10^3^ PFU. **f.** Schematic showing proposed mechanism of øAbs1 infectivity across *A. baumannii* strain D86 wild-type, Δ*galU*, Δ*oprC*, and Δ*galU:ΔoprC* double mutant. Black arrow indicates infection, blue flat line indicates no infection. **g.** Growth curves (OD_600_) of *A. baumannii* D86 mutants represented in panel f. Bacteria only (black) and in the presence of øAbs1 (red) at an MOI of 1. Data are median ± 95% CI (n = 9). **h.** Scanning electron microscopy (SEM) images of *A. baumannii* D86 mutants represented in panel f, all were infected with øAbs1 with a minimum of 100 cells imaged per condition. White arrows indicate adsorbed virions (scale bar = 500 nm).

Working on this assumption, we tested øAbs1 infectivity against four *A. baumannii* strains, spanning different KLs that include both wild-type and CPS-deficient mutants with decreased production of CPS (**Fig. 1d**). We found that øAbs1 infection correlates with decreased production of CPS, except for its host of isolation strain LUH5538 (**Fig. 1e**). Given our prior attempts to identify the receptor of øAbs1 failed, we adopted an alternative strategy using the decapsulated mutant *A. baumannii* strain D86 (KL3v1) *galU_C652T*, which had a verified loss-of-function mutation in its *galU* gene (stop gained, C652T > Gln218*); a critical enzyme for synthesizing sugar precursors required to build the CPS^5^. We performed multiple rounds of directed evolution using øAbs1 and *A. baumannii* strain D86 *galU_C652T* to isolate a stable øAbs1-resistant mutant. Whole-genomic sequencing and analysis confirmed the presence of the prior *galU* mutation, along with a four-nucleotide insertion in the *oprC* gene (insertion, 1489_1494insGACC) leading to a frameshift and a truncated protein of 519 amino acids, compared with the original protein of 704 amino acids. To validate the *oprC* gene product as the phage receptor for øAbs1, we generated single knockout mutants of *galU* and *oprC*, as well as a double knockout lacking both genes in *A. baumannii* strain D86, all of which were confirmed via PCR and Sanger sequencing. We then evaluated øAbs1 infectivity against wild-type D86 and its knockout variants (**Fig. 1f**). In the presence of øAbs1, we observed unimpeded growth of D86 wild-type, Δ*oprC* single knockout, and the Δ*galU*:Δ*oprC* double knockout, suggesting there was no infection in these mutants (**Fig. 1g**). In contrast, the Δ*galU* mutant, which lacks CPS but contains OprC, was highly susceptible to øAbs1 infection, with a sustained antibacterial effect lasting over 24 h. To supplement this, we performed scanning electron microscopy (SEM) across these phage-host pairs, observing øAbs1 adsorption to the Δ*galU* mutant and no attachment to the other strains (**Fig. 1h**, minimum 100 cells observed per condition).

The *oprC* gene codes for a TonB-dependent outer membrane transporter responsible for copper homeostasis and bacterial virulence^12,13^. Given the importance of copper transport during *A. baumannii* infection^13^, we undertook a bioinformatic survey to determine the prevalence of *oprC* in 165 clinical *A. baumannii* isolate genomes (Extended Data Table 1)^14,15^. We found 86% of the genomes (142/165) harboured a complete gene copy of *oprC* (**Fig. 2a**; >90% coverage, >95% amino acid identity) with an average protein length of 698 amino acids (s.d. ±50 amino acids), which was consistent with the expected size for TonB-dependent outer membrane receptors^13^. To assess redundancy and clustering, all 142 OprC-like proteins were dereplicated at 100% amino acid identity across the full length, yielding 53 distinct clusters (Extended Data Fig. 1b), with most proteins belonging to large clusters with high internal similarity. We then experimentally confirmed øAbs1 infection in ten CPS-deficient *A. baumannii* mutant strains, spanning KL2, KL3, KL11, KL14, and KL22 in our collection, all of which contained a complete gene copy of *oprC* (**Fig. 2a**; Extended Data Table 1). This suggests that *oprC* is evolutionarily conserved across clinical *A. baumannii* strains, despite extensive variation in other surface structures (i.e. KL), which often diversify rapidly under phage selection pressures^16^.

**Figure 2.**
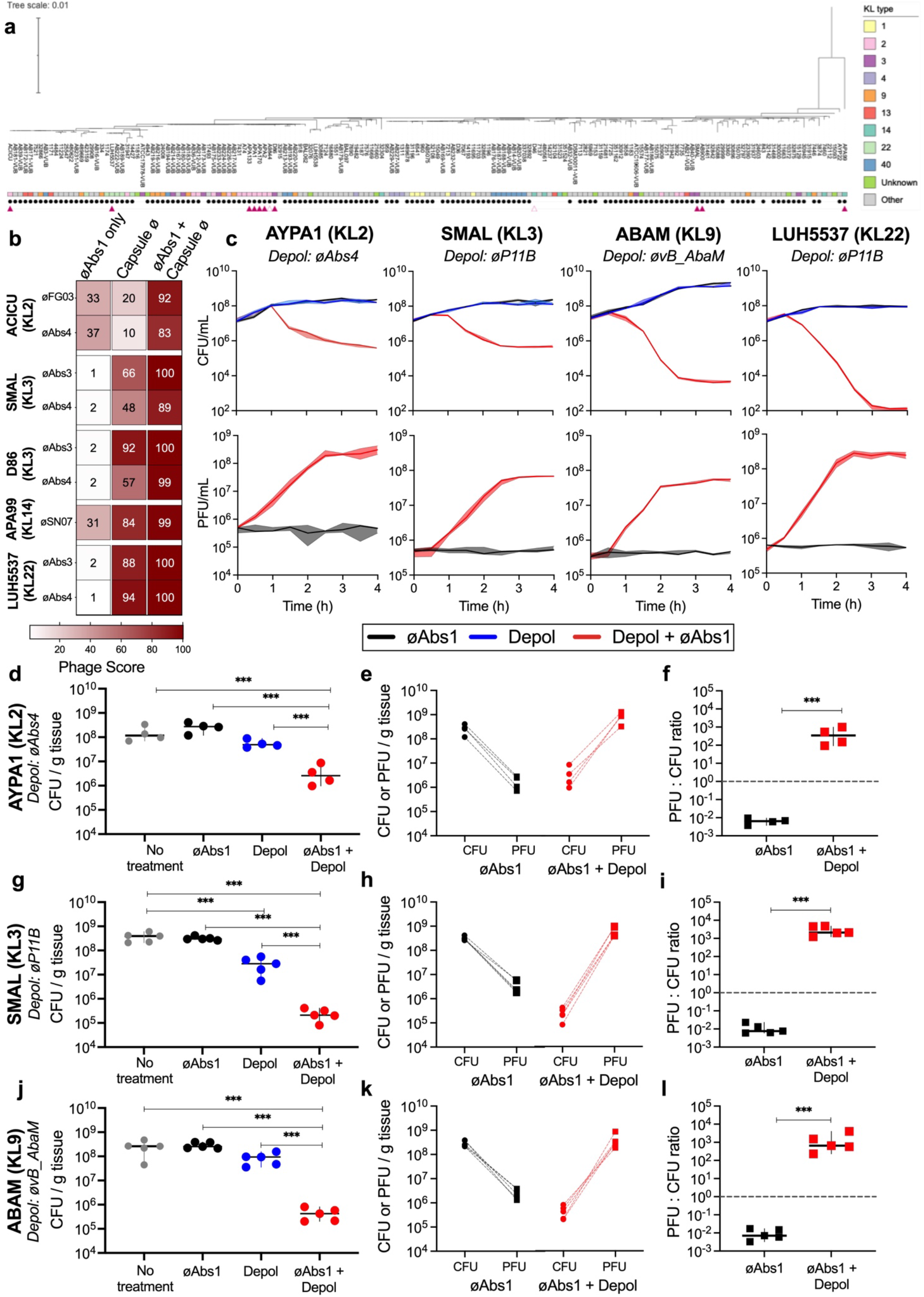
**a.** Phylogenetic tree of 165 clinical isolates of *A. baumannii* showing the prevalence of OprC. Solid rectangles denote the KL type, black circles indicate isolates with a complete gene copy of *oprC* (>90% coverage, >95% amino acid identity), solid red triangles indicate isolates with confirmed øAbs1 infectivity when capsule was disrupted, empty red triangles isolates lacking øAbs1 infectivity. **b.** Heat map showing infectivity of øAbs1 alone, a collection of capsule-targeting phages (øAbs3, øFG03, øAbs4, and øSN07) alone, and their combination, across five clinical *A. baumannii* strains of diverse KL types. Heat map represents Phage Score^23^, which takes growth kinetic data and integrates these into a value from 0 to 1 (shown here on 100-point scale), with higher values (red) representing greater phage infectivity. Data are mean (n = 9). **c.** Time course infection assays over four hours showing Colony Forming Units (CFU/mL; top) and Plaque Forming Units (PFU/mL; bottom) of four *A. baumannii* strains from diverse KL types grown in the presence of øAbs1 (black), a matched depolymerase (blue), or a combination of øAbs1 and matched depolymerase (red). Data are mean ± 95% CI (n = 3). **d-f.** Paired data from murine bacteraemia model of BALB/c mice infected with clinical *A. baumannii* strain AYPA1 (KL2). Panel d; averaged bacterial load across four respective treatments; no treatment control (gray), øAbs1 only (black), depolymerase only (blue), and combination of øAbs1 and depolymerase (red). Each symbol represents average bacterial burden (CFU/g tissue) from four body sites of an individual mouse, line shows geometric mean ± 95% CI (n = 4), statistical test one-way ANOVA, on data as log_10_ CFU followed by Tukey’s test for all pairs; *** p<0.001. Panel e; paired CFU and PFU measures from mice treated with øAbs1 alone (black) or combination of øAbs1 and depolymerase (red). Each symbol represents average bacterial or phage counts from four body sites of an individual mouse (n = 4). Panel f; ratio of PFU to CFU from mice treated with øAbs1 alone (black) or combination of øAbs1 and depolymerase (red). Each symbol represents average bacterial or phage counts from four body sites of an individual mouse (n = 4), comparisons with Welch’s t-test on data as log_10_ values; *** p<0.001. **g-i.** Paired data from murine bacteraemia model with clinical *A. baumannii* strain SMAL (KL3). Panels g, h, and i, ordered and described as above, (n = 5). **j-l.** Paired data from murine bacteraemia model with clinical *A. baumannii* strain ABAM (KL9). Panels j, k, and l, ordered and described as above (n = 5).

Given that OprC is a prevalent and highly conserved outer membrane protein that acts as the phage receptor for øAbs1, we reasoned that provided the CPS was degraded to expose OprC, øAbs1 could function as a broad-spectrum therapeutic against clinical *A. baumannii* strains. As a proof of concept, we examined bacterial growth in liquid culture over 24 hours in the presence of øAbs1, one of four CPS-targeting phages (øAbs3, øFG03, øAbs4, and øSN07), or their combination to test for combined effects. We tested these phages across five prevalent *A. baumannii* KL types (strains ACICU [KL2], SMAL [KL3], D86 [KL3v1], APA99 [KL14], and LUH5537 [KL22]), all of which carried the *oprC* gene (**Fig. 2b**). Capsule-targeting phages on their own had moderate-to-high bacterial suppression. Comparatively, øAbs1 on its own had little-to-no effect on the capsulated strains, yet when combined with capsule-targeting phages, resulted in strong bacterial suppression, often without emergence of phage-resistance (Extended Data Fig. 2).

Due to the complexity of multi-phage co-infection assays, we theorized that phage-encoded depolymerases, which are specialized enzymes encoded on phage tail fibres that recognize and degrade bacterial CPS to facilitate infection^17^, should function to remove bacterial capsule and enable øAbs1 infection via the OprC receptor. We obtained the genome sequences of three phages (øAbs4, øPB11^18^, øvB_AbaM^19^), which infected strains displaying the four most prevalent *A. baumannii* KL types^8^ (AYPA1 [KL2], SMAL [KL3], LUH5537 [KL22], and ABAM [KL9]). From these phage genomes, we identified candidate depolymerase genes using DepoScope^20^ (Extended Data Table 2), which were subsequently expressed in *Escherichia coli* with recombinant depolymerases purified via affinity chromatography. We then combined øAbs1 with each depolymerase individually and quantified both bacterial killing and phage replication alongside respective controls (**Fig. 2c**). In all tested strains, the combination of øAbs1 and depolymerase reduced bacterial growth and enabled øAbs1 replication in capsulated isolates that did not support øAbs1 replication alone.

Finally, we sought to evaluate the therapeutic potential of øAbs1 and depolymerases within a pre-clinical *in vivo* model. Using our established murine model of bacteremia^21^, groups of BALB/c mice were given single intraperitoneal (IP) injection of a clinical *A. baumannii* strain (1×10^6^ CFU) from three KL types (AYPA1 [KL2], SMAL [KL3], and ABAM [KL9]) to establish acute bacteraemia. Mice were randomized into four treatments groups (four-to-five mice per group): no-treatment control, øAbs1 only (1×10^6^ PFU, MOI = 1), depolymerase only (1.65 µM), and combination of øAbs1 with depolymerase against each specific KL type (1×10^6^ PFU, 1.65 µM). One hour-post-infection (hpi), mice were given a single IP dose of the respective treatment, and eight to nine hpi, were euthanized with bacterial and phage counts assessed (**Fig. 2d-l**; see Extended Data Fig. 3 for per body site counts). Across the three strains, the combination of øAbs1 and depolymerase reduced bacterial burden by 50- to 1,600-fold relative to untreated mice, whereas øAbs1 alone had no significant effect, and depolymerase alone reduced CFU significantly in only one strain (SMAL [KL3]). Further, in the presence of depolymerase, øAbs1 titres were 510-fold higher, while bacterial burden was 93-fold lower, such that phage outnumbered bacteria in every mouse receiving the combination treatment. Taken together, these results support depolymerases helping øAbs1 to reach and replicate in *A. baumannii* hosts it otherwise cannot infect.

In conclusion, we characterize the virulent øAbs1 that infects decapsulated *A. baumannii* strains via the TonB-dependent receptor OprC, which is highly conserved in clinical isolates. We demonstrate that a mechanistic knowledge of phage receptor biology and infectivity^22^, enabled a combinatorial strategy of phage-encoded depolymerases with øAbs1 to target and kill this critical-priority pathogen. Importantly, this strategy overcomes a major barrier for the implementation of phage therapy to treat *A. baumannii* infections, being the limited phage host range coupled with extensive CPS diversity. In summary, the combination of phage Abs1, which targets a highly conserved and functionally important non-capsular bacterial receptor, alongside recombinant depolymerases that degrade *A. baumannii* capsular polysaccharides, could enable a broad-spectrum antimicrobial treatment against this critical priority pathogen.

## ONLINE METHODS

### Bacterial strains and plasmids

Extended Data Table 3 contains a list of bacterial strains and phages used in this study

### Phage isolation and purification

Phages were isolated from raw sewage samples, kindly provided by colleagues in the Monash University Engineering Department. Aliquots of centrifuged and 0.22 µm filtered sewage were combined with overnight cultures of *A. baumannii* in lysogeny broth (LB) and incubated overnight at 37 °C with shaking. Resulting lysates were purified following the previously described Phage-on-Tap protocol^24^, and screened for phage presence using the standard double overlay agar assay.

### Phage DNA extraction and sequencing

Phage lysates were pretreated with DNase I and RNase A (1 µg/mL each) for 2 hours at 37 °C, followed by enzyme inactivation at 75 °C for 5 minutes. Next, 1.8 mL aliquots were mixed with 92 µL of 10% (w/v) SDS, 18 µL of proteinase K (10 mg/mL), and 5 µL of glycogen (20 mg/mL) were added and incubated at 37 °C for 30 minutes. Lysates were added to phase-lock gel tubes and extracted with equal volumes of phenol:chloroform:isomyl alcohol (25:24:1, v/v/v; pH 6.7) and separated by centrifugation at 1500 rpm for 5 minutes. The upper aqueous phase was transferred to a new phase-lock gel tube and extraction repeated, followed by a final purification step using 500 µL of chloroform:isoamyl alcohol (24:1, v/v), followed by centrifugation at 6000 rpm for 5 minutes. Aqueous phase was transferred to a new tube and DNA precipitated by addition of 45 µL of 3 M sodium acetate (pH 5.2) and 500 µL of ice-cold 100% isopropanol, followed by incubation at room temperature for 20 minutes. The precipitate was pelleted by centrifugation at 16,000 rpm for 20 minutes at 4 °C, supernatant discarded, and DNA pellet washed twice with 1 mL of 70% ethanol. The dried DNA pellet was resuspended in 200 µL TE buffer (10 mM Tris-HCl, 1 mM EDTA, pH 8.0). The integrity of extracted genomic DNA was evaluated by electrophoresis on 0.7% (w/v) agarose gels prepared in Tris-Acetate-EDTA (TAE) buffer stained with GelRed^®^ Nucleic Acid Stain (Merck). Qualified samples were submitted for library preparation and next-generation sequencing using the Illumina HiSeq 150 bp paired-end platform (Genewiz, China).

### Transmission electron microscopy (TEM)

Phage suspensions were deposited onto a copper TEM grid (200 mesh, SPI Supplies) precoated with a carbon-coated ultrathin formvar film, incubated for 30 seconds, and blotted dry with filter paper. The grids were then stained by applying 10 µL of 1% (w/v) uranyl acetate-water solution to the grid surface for 20 seconds, followed by drying with filter paper and examined using a JEM-1400 Plus (Jeol) operated at an accelerating voltage of 80 kV.

### Growth curve assay

Bacterial growth kinetics were monitored by measuring OD_600_ in a microplate reader (Epoch Microplate Spectrophotometer; BioTek, Winooski, VT, USA). Exponentially growing bacterial cultures were added to individual wells of a 96-well plate and combined with phage lysates at a predetermined multiplicity of infection (MOI). Plates were incubated at 37 °C with continuous shaking, and absorbance readings were recorded every 15 minutes for up to 24 hours. Phage activity was quantified by calculating a phage score ^23^, based on the area under curve (AUC) of bacterial growth in the presence and absence of phage.

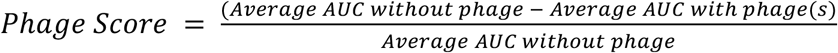

### One-step growth curve

Overnight bacterial cultures were diluted to an OD_600_ of 0.3 (∼5 × 10^7^ colony forming units [CFU]/mL) and checked by viable cell counts. Phage was added at a multiplicity of infection (MOI) of 0.01 (∼5 × 10^5^ plaque forming units [PFU]/mL) and incubated at 37 °C with shaking. After 2 minutes, cultures were centrifuged (12,000 rpm, 1 minute) to remove unadsorbed phage, the pellets were resuspended in fresh medium, diluted 1:1,000, and incubated at 37 °C with shaking. Samples were taken in triplicate at defined time points and titrated using top-agar overlays. Burst size was calculated by dividing the PFU at the first post-lysis plateau by the PFU during the latent period.

### Capsule quantification using phenol-sulfuric acid method

Bacterial CPS was quantified using the phenol-sulfuric acid method^25^. Bacterial cultures were standardized to a fixed cell number per sample (2 × 10^9^ CFU), determined by OD_600_ measurements and confirmed via viable cell counts. Cell suspensions were treated with phenol and sulfuric acid to hydrolyse and quantify polysaccharides based on colorimetric detection at 490 nm (OD_490_). Calibration curves were generated using glucose standards to ensure accurate quantification.

### Generation of resistant mutants and variant calling

Phage-resistant mutants were generated by exposing bacterial cultures to their respective phages for three consecutive daily passages, with centrifugation and replacement of supernatant between passages to prevent phage co-evolution. Resistance was confirmed via soft-agar overlay and growth curve assays. Bacterial genomic DNA was extracted using the GenElute™ Bacterial Genomic DNA Kit (Sigma-Aldrich, Australia), quantified, assessed for purity, and vacuum dried for transport. Sequencing was carried out on the Illumina HiSeq platform (Genewiz, China). Single-nucleotide variants (SNVs) and small insertions/deletions were identified using Snippy v4.3.6 (https://github.com/tseemann/snippy) with default parameters, using the assembled genome of wild-type as the reference assembly.

### Generation of knockout mutants and validation

Knockout mutants of *A. baumannii* were constructed using an allelic exchange approach based on previously described protocols employing suicide vectors^26^, with minor modifications tailored to our strains. Approximately 1,000 bp of DNA flanking both the upstream and downstream regions of the target gene were synthesised to form the linear knockout fragment. The resulting fragment was cloned into a pALFI3 allelic exchange vector and transformed into *E. coli* Jke201 for biparental mating. Donor (*E. coli* Jke201) and recipient *A. baumannii* strains were streaked on LB agar supplemented with 50 µg/mL ampicillin and 100 µM diaminopimelic acid (DAP). The following day, 25 µL of donor and recipient suspensions were mixed and spotted onto a dry LB + DAP (100 µg/mL) plate. The next day, a loopful of cells was taken from the conjugation patch was resuspended in 100 µL LB broth and plated onto selective agar containing 50 µg/mL tellurite. Putative single-crossover mutants were re-streaked on selective media containing 50 µg/mL tellurite and verified by colony PCR. These single-crossover mutants were then streaked onto sucrose-based counterselection plates composed of zero-salt LB medium (10 g/L tryptone, 5 g/L yeast extract, 100 g/L sucrose, 15 g/L agar) supplemented with 10% sucrose (w/v). Individual colonies were patched onto LB plates containing tellurite (50 µg/mL) and LB + 10% sucrose (w/v) plates to identify colonies that had lost the vector backbone. Colonies that grew on sucrose but not on tellurite plates represent potential successful mutants, and were subsequently verified by colony PCR and Sanger sequencing to confirm the loss of the target gene.

### Scanning electron microscopy (SEM)

A 20 µL bacterial suspension (∼1 × 10^7^ colony-forming unit (CFU/mL) was deposited onto a fresh gold-coated silicon wafer (5 × 5 mm) and cells were allowed to set for 5 minutes. The wafer was briefly washed in LB medium and then placed (face side up) on filter paper. 20 µL of phage lysate (∼1 × 10^8^ PFU/mL) was then applied and allowed to incubate for 20 minutes, then the wafer was intensively washed in PBS, and then immersed in 2.5% glutaraldehyde in PBS for 15 minutes, washed, and progressively dehydrated through increasing concentrations of ethanol. After processing in a critical point dryer Leica CPD300, the prepared wafer was mounted on a standard aluminium 12.7 mm SEM stub and coated with a ∼10 nm thick gold layer with a BAL-TEC AG sputter coater SCD 005. Imaging was conducted using the FEG-SEM ThermoFisher Helios 5 Hydra at an accelerating voltage of 2 kV in secondary electron mode, with a 2 mm working distance, operating in immersion mode with the through-the-lens detector.

### Bacterial taxonomic annotation and phylogenetic tree construction

Predicted proteomes of *Acinetobacter baumannii* were collected from two publicly available GenBank Bioprojects PRJNA798866^15^ and PRJNA545079^14^, and the predicted proteome from our local collection of *A. baumannii* isolates. Proteins homologous to the reference *A. baumannii* strain D86 OprC protein were searched using BLAST+ v2.9.0^27^. Homology was detected when the target protein covered at least 90% of the reference OprC with at least 95% of amino acid identity. An additional search using HMMER 3.4 was performed^28^, but no additional homologue was detected. To prepare the protein dataset for phylogenetic reconstruction, identical homologous OprC proteins were dereplicated based on a local alignment using CD-HIT v4.8.1 (-c 1 -n 5 -g 1 -d 0 -G 0 -aS 1)^29^, and the protein with the longest amino acid sequence was established as the representative protein of each cluster. Representative OprC amino-acid sequences were aligned with MAFFT v7.525 (default settings)^30^ and the alignment was cleaned using CIAlign (default settings) to remove poorly aligned or gap-rich regions^31^; the cleaned alignment was then used to infer a maximum-likelihood protein phylogeny with IQ-TREE v2.0-rc1 (-m MFP -bb 1000)^32^, for which JTT+I+G4 was identified as the best-fit protein substitution model based on Bayesian Information Criterion, and the resulting tree was visualized in Interactive Tree of Life (iTOL)^33^. To generate a whole-genome phylogeny, each genome was taxonomically assigned according to the Genome Taxonomy Database (GTDB) (release 220) using GTDB-Tk v2.4.0 (--write_single_copy_genes --keep_intermediates --skip_ani_screen)^34^. The protein sequence alignment of 120 core bacterial genes generated by GTDB-Tk was then employed to build a phylogenetic tree using IQ-TREE v.2.0-rc1 (-m MFP -bb 1000)^32^, for which mtVer+F+R2 was identified as the best-fit protein substitution model based on Bayesian Information Criterion, and this tree was likewise visualised and annotated in iTOL to permit mapping of OprC presence/absence and other metadata onto the species phylogeny^33^.

### Cloning and purification of phage depolymerases

Depolymerase genes were identified from the genomes of capsule-targeting phages ΦAbs4 (this study), PB11^18^, vB_AbaM_IME285^19^ using DepoScope^20^ (Extended Data Table 2). Full-length coding sequences of the identified genes were subsequently synthesized by Integrated DNA Technologies. The sequences were cloned into the modified pET-28a vector encoding an N-terminal His_6_ tag, with or without a Small Ubiquitin-like Modifier (SUMO) tag. The resulting plasmids were transformed into *Escherichia coli* BL21(DE3). Bacterial cultures were grown in LB medium supplemented with 30 μg/mL kanamycin at 37 °C until an OD_600_ of 0.6 - 0.8 was reached. The protein expression was induced by adding isopropyl β-D-1-thiogalactopyranoside (IPTG) to a final concentration of 0.5 mM. Cells were harvested by centrifugation, resuspended in lysis buffer (20 mM Tris pH 8.0, 500 mM NaCl) and disrupted by sonication to release intracellular contents. Lysates were clarified by centrifugation and depolymerases were purified from the soluble fraction using immobilised metal affinity chromatography (5 ml His-Trap HP nickel column, Cytiva). If present, the SUMO tag was subsequently cleaved and removed, followed by size-exclusion chromatography using a Superdex 200 column.

### Murine model of *A. baumannii* bacteraemia

The murine bacteraemia model employed in this study was largely adapted from previously established protocols^9,21^.In brief, experiments were conducted using female BALB/c mice, six to ten weeks of age and weighing at least 15 g, with each animal serving as a single experimental unit. Mice were randomly assigned to treatment groups to minimise bias. Bacterial inoculums were prepared from late-exponential phase *A. baumannii* cultures and adjusted to 1 × 10^6^ CFU. At this dose, untreated mice consistently reached the clinical score threshold for humane euthanasia by nine hours post-infection. To enhance virulence, the inoculum (100 μL in phosphate-buffered saline (PBS)) was mixed 1:1 with 6% porcine stomach mucin (Sigma-Aldrich, dissolved in PBS)^36^, yielding a final injection volume of 200 μL. All infections and treatments were administered intraperitoneally. Treatment volumes were standardised to 200 μL per mouse, with PBS used in place of active treatment when necessary to ensure uniform injection volumes across groups and time points. At experimental endpoints, mice were euthanised via CO₂ asphyxiation. Blood and organs (liver, spleen, and kidney) were harvested, weighed, and homogenised in PBS. Homogenates and blood samples were serially diluted and plated for CFU enumeration, normalised to organ weight, which served as the primary outcome measure. For mice that received phage treatment, homogenates and blood suspensions were centrifuged, and the supernatants were serially diluted and assessed by soft agar overlay for PFU quantification.

## Ethics statement

All protocols involving animals were reviewed and approved by the Monash University Animal Ethics Committee (Project ID: 45807), and the animals were housed at the Monash Animal Research Platform, Monash University, under standard laboratory conditions: 20-22 °C, 30-70% relative humidity, and 12:12 hour light/dark cycle. We conducted all procedures in strict adherence to institutional guidelines.

## Data representation and statistical analyses

All graphing and statistical analyses were conducted using GraphPad Prism version 11.0.1 (GraphPad Software, Inc.). Schematics were created with BioRender.com. For animal experiments involving four groups, CFU and PFU values were first transformed using the formula log_10_(Y+1) to normalize variance. Normality was assessed using the Shapiro-Wilk test, supplemented by a visual inspection of QQ-plots. Homogeneity of variance was evaluated using Brown-Forsythe’s test, with additional confirmation by visual inspection of residual plots to detect potential funnel-shaped patterns, which can indicate heteroscedasticity. When data failed the Shapiro-Wilk test, indicating a non-Gaussian distribution, the non-parametric Kruskal-Wallis ANOVA was applied. When variance was unequal despite normality (i.e., Shapiro-Wilk passed but Brown-Forsythe or residual plots suggested heteroscedasticity), Welch’s ANOVA was performed. If both assumptions were satisfied, a standard one-way ANOVA was used. Post hoc analyses for each sample category were selected according to biological relevance. Specifically, comparisons were primarily focused on the phage combination group versus each individual treatment, with multiple comparisons corrected using Dunnett’s T3 test. For comparisons involving all groups across averaged sample types, the Tukey’s multiple comparisons test was applied. For animal experiments involving two groups, the same data transformation (log_10_(Y+1)) and normality checks (Shapiro-Wilk test and QQ-plots) were performed. Equality of variance was assessed using F test, again supported by residual plot inspection. If data were non-Gaussian (Shapiro-Wilk failed), the Mann-Whitney U test was applied. If variances were unequal (F test failed or residual plots indicated heteroscedasticity), Welch’s corrected unpaired t-test was used. If both assumptions were met, a standard unpaired t-test was performed.

## Code Availability

The study did not involve the creation of novel code for bioinformatics analyses. All the used software is freely accessible and is referenced.

## Acknowledgements

This work, including the efforts of J.J.B., was funded by the National Health and Medical Research Council (NHMRC) Investigator Award (2026130). We acknowledge the Monash Animal Research Facility (MARP) for providing the necessary facilities for animal experiments; the MASSIVE HPC facility (www.massive.org.au) for providing us with cluster time for data analysis; and the Monash Ramaciotti Cryo-EM platform for the use of facilities.

## Author contributions

LK and JJB conceived the study. LK performed all *in vitro* experimental work. LAF and FGA contributed to the bioinformatic analyses. LK, DS, SRD, SK, and RB carried out depolymerase identification and cloning. LK, DS, and MC conducted *in vivo* infection experiments. IMB and EG assisted with phage genome extraction and assembly. LK and DK performed the electron microscopy experiments. RMH, JJK, and AYP provided access to clinical strains. RMH and JJK provided additional investigation and characterisation of clinical strains and KL types. JJB, FGA, and AYP contributed to supervision. JJB led and funded the study. LK and JJB wrote the original draft. All authors reviewed and approved the final manuscript.

## Extended Data

**Extended Data Figure 1.**
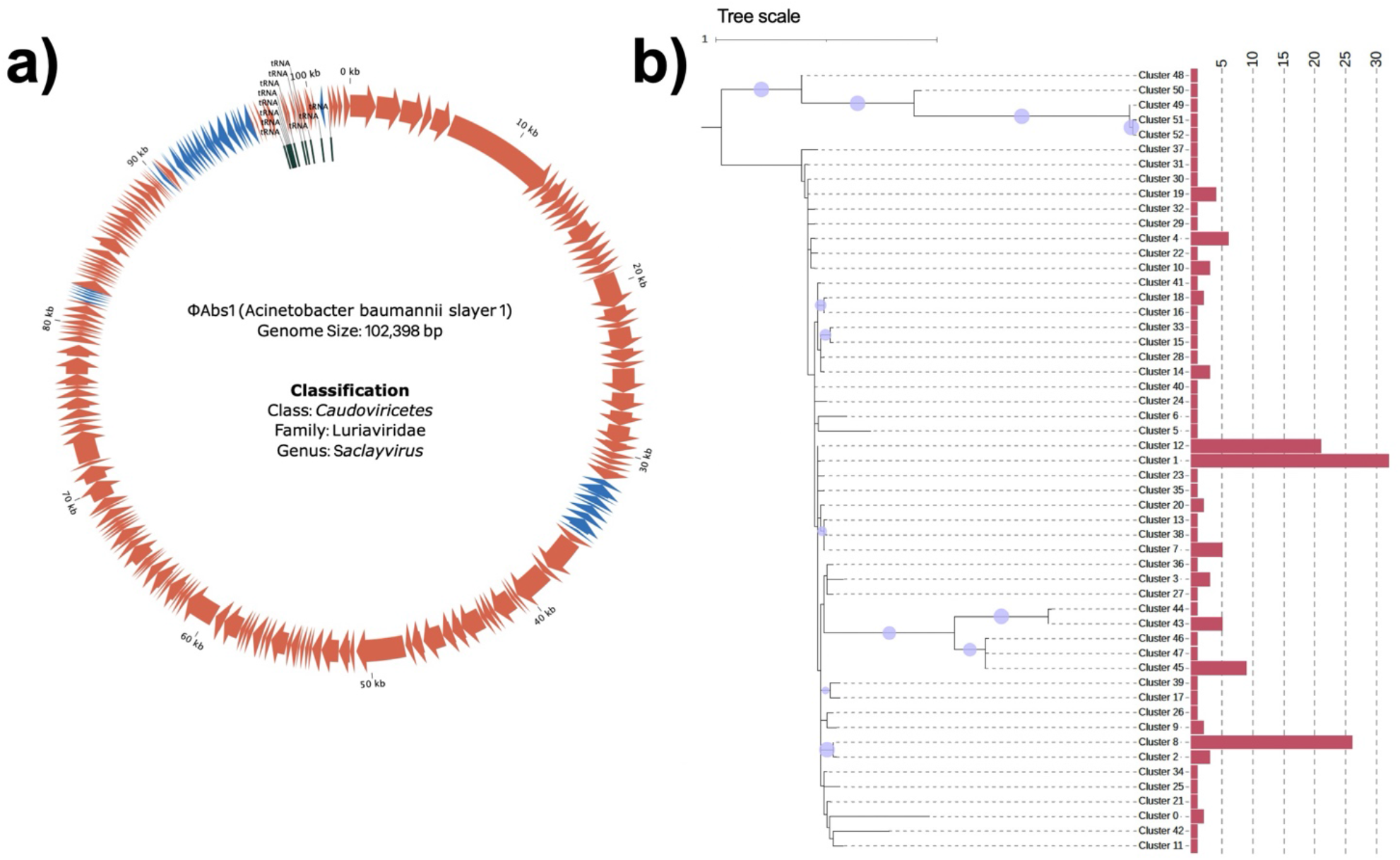
**a)** Genome of øAbs1 (102,398 bp) that was classified as Order *Caudoviricetes*, Family *Luriaviridae*, Genus *Saclayvirus*. Orange arrows indicate ORFs in forward direction and blue arrows indicate ORFs in reverse direction. **b)** Consensus phylogenetic tree of OprC amino acid clusters. The 142 OprC protein sequences were dereplicated at 100% identity across 100% of shorter sequence length. The number of proteins within each cluster is indicated by pink bars. The tree is rooted at the midpoint, and internal dots denote boostrap values greater than 95%. The reference *A. baumannii* strain D86 is in cluster 1.

**Extended Data Figure 2.**
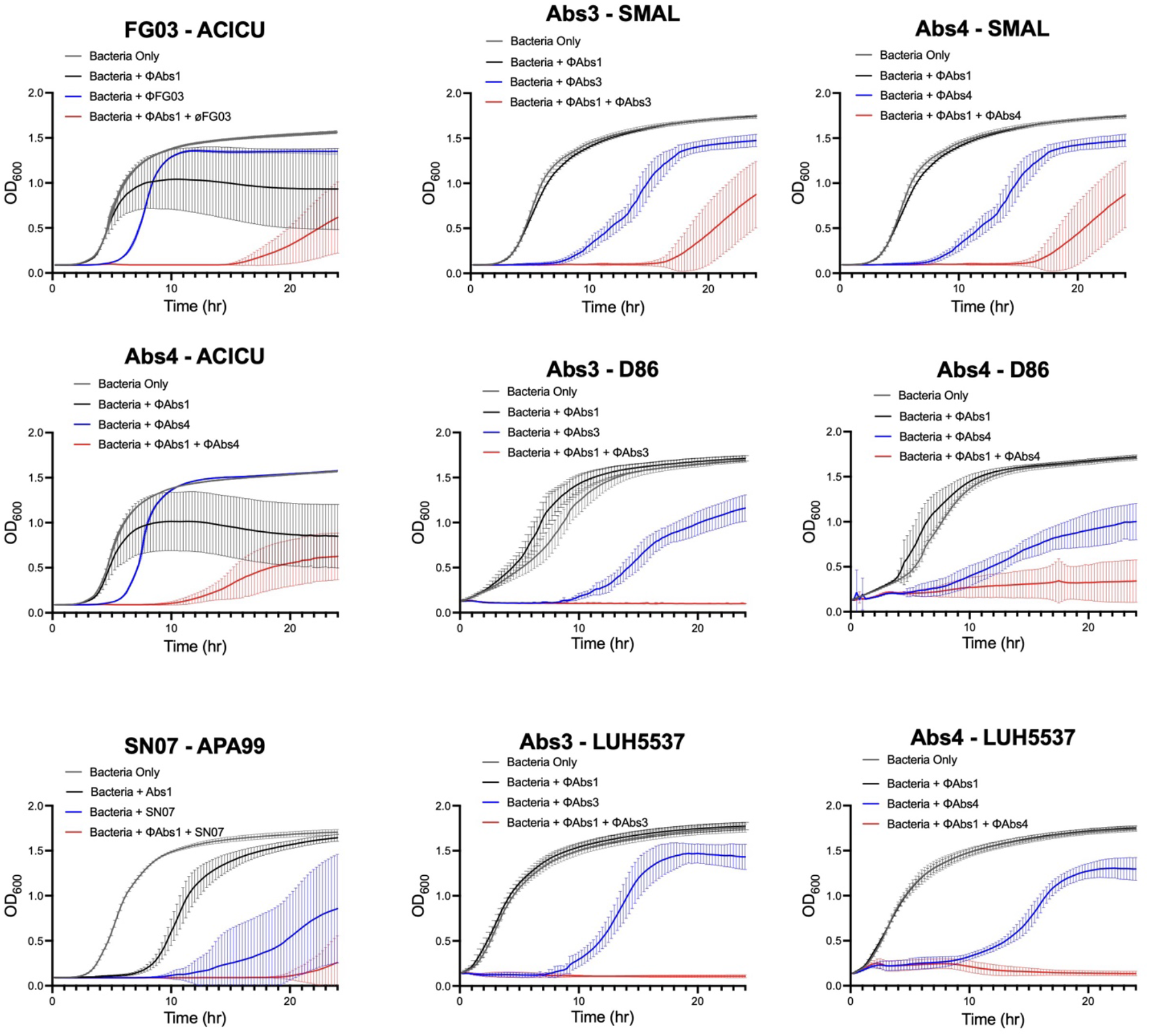
Individual growth curves of *A. baumannii* host strains on their own (gray), in the presence of øAbs1 (black), in the presence of a capsule-targeting phage (blue), and in combination with øAbs1 and capsule-targeting phage (red). All experiments at MOI of 1. Line are median ± 95% CI (n = 9). OD_600_, optical density at 600 nm.

**Extended Data Figure 3.**
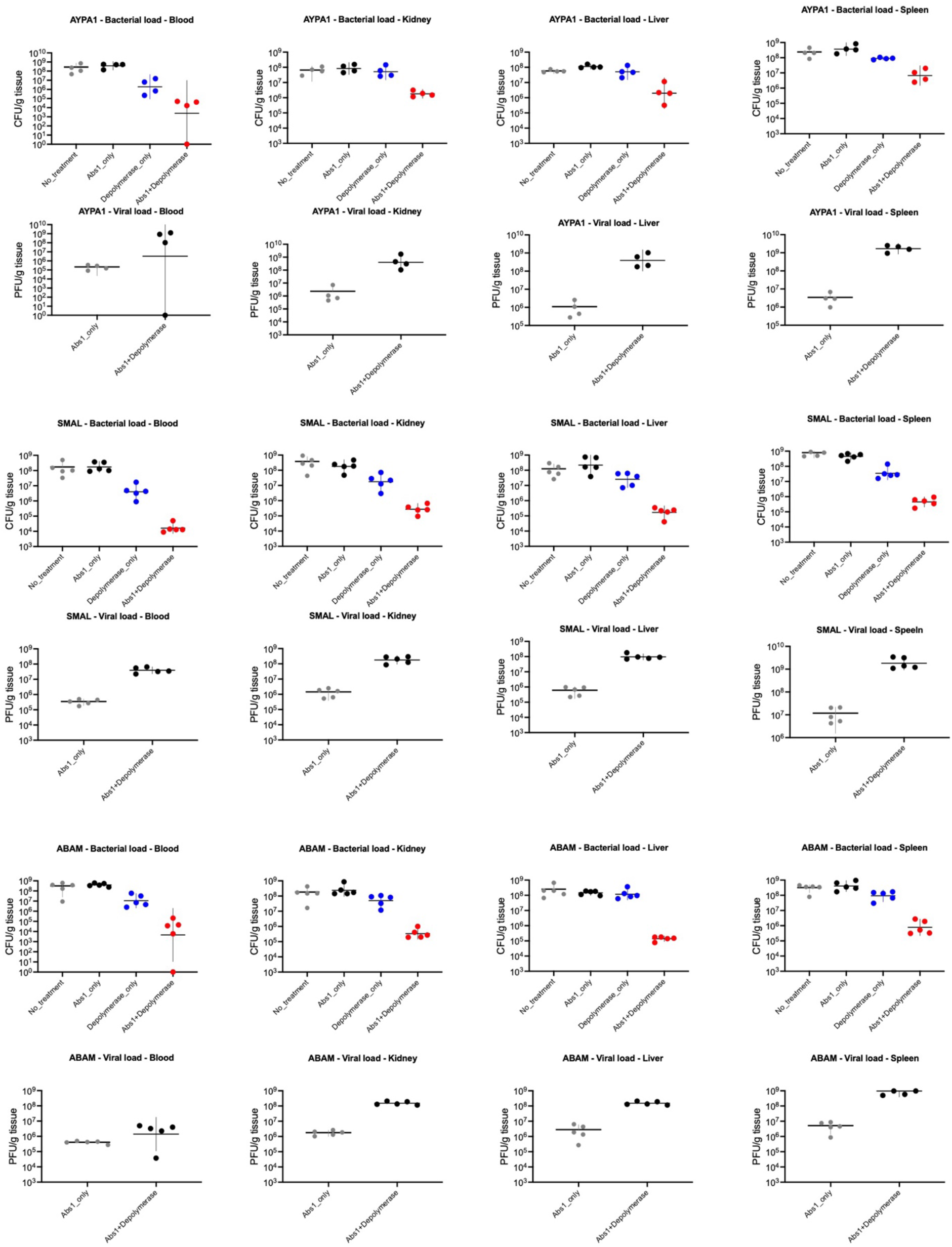
Individual CFU and PFU counts per gram of tissue from murine bacteraemia model of BALB/c mice infected with clinical *A. baumannii* strains. Each symbol represents average bacterial (CFU/g tissue) or phage (PFU/g tissue) from each body sites of an individual mouse, line shows geometric mean ± s.d. (n = 4-5).

**Extended Data Table 1.** Source information for Acinetobacter baumannii genomes used in Figure 2a. 165 clinical *A. baumannii* isolate genomes obtained from both public datasets (n = 146)^14,15^ and our in-house culture collection (n = 19).

**Extended Data Table 2.** Depolymerase sequences cloned from phage genomes.

**Extended Data Table 3.** List of bacterial strains and phages used in this study.

